# Effects of the G-quadruplex-binding drugs Quarfloxin and CX-5461 on the malaria parasite *Plasmodium falciparum*

**DOI:** 10.1101/2023.07.13.548827

**Authors:** Holly M. Craven, Guilherme Nettesheim, Pietro Cicuta, Andrew M. Blagborough, Catherine J. Merrick

## Abstract

*Plasmodium falciparum* is the deadliest causative agent of human malaria. This parasite has historically developed resistance to many drugs, including the current frontline treatments, so new therapeutic targets are needed. Our previous work on guanine quadruplexes (G4s) in the parasite’s DNA and RNA has highlighted their influence on parasite biology, and revealed G4 stabilising compounds as promising candidates for drug repositioning. In particular, quarfloxin, a former anticancer agent, kills blood-stage parasites at all developmental stages, with fast rates of kill and nanomolar potency. Here we explored the molecular mechanism of quarfloxin and its related derivative CX-5461. *In vitro,* both compounds bound to *P. falciparum*-encoded G4 sequences. *In cellulo*, quarfloxin was more potent than CX-5461, and could prevent establishment of blood-stage malaria *in vivo* in a murine model. CX-5461 showed clear DNA damaging activity, as reported in human cells, while quarfloxin caused weaker signatures of DNA damage. Both compounds caused transcriptional dysregulation in the parasite, but the affected genes were largely different, again suggesting different modes of action. Therefore, CX-5461 may act primarily as a DNA damaging agent in both *Plasmodium* parasites and mammalian cells, whereas the complete antimalarial mode of action of quarfloxin may be parasite-specific and remains somewhat elusive.

## Introduction

Malaria is a disease caused by infection with parasitic apicomplexan *Plasmodium* species, transmitted through bites of female *Anopheles* mosquitoes. The disease is widespread throughout the global south and causes significant morbidity and over half a million deaths annually [1]. Of the six species known to cause human malaria, *Plasmodium falciparum* causes the most deaths. *P. falciparum* is endemic throughout sub–Saharan Africa and much of Asia, and the disease is a major barrier to socio-economic development. Despite being managed in many parts of the world through a rigorous programme of insecticide spraying, insecticide-treated bed net distribution and antimalarial treatment, resistance to both insecticides and drugs commonly arises [2–6] and can spread rapidly [7]. Although current programmes of artemisinin-based combination therapy (ACT) have greatly reduced the clinical burden of malaria, more new drug targets are needed.

Here, we focus on the guanine quadruplex (G4) as a potential antimalarial target. Being a nucleic acid structure rather than a protein, the G4 would represent an entirely novel drug target in the parasite. *P. falciparum* has a highly A/T-biased genome (∼81% A/T) with very few sequences that have the potential to form G4s (‘PQSs’); nevertheless, we and others have shown that these G4s do exist and that they play roles in *Plasmodium* biology at both the DNA [8–11] and RNA [12,13] levels. The existence of G4s in malaria parasites raises the possibility of repositioning G4-binding drugs – which are often developed primarily as anticancer agents – as antimalarial agents. Drug repositioning can be a valuable strategy for tropical disease treatment, due to limited funds for drug development [14–16].

We previously showed that a G4-binding agent called quarfloxin, which previously reached phase IIa clinical trials for cancer [17–21], is potent and fast-acting against *P. falciparum* parasites cultured *in vitro* [10]. However, the drug did not act mechanistically via the route characterised in human cells, and subsequently also in trypanosomes [20–23], i.e. disrupting rRNA production by RNA polymerase I. This was perhaps unsurprising, since rRNA genes in *Plasmodium* do not occur in the characteristic tandem arrays found in many eukaryotes [24]. Importantly, this suggested that quarfloxin may have a parasite-specific mode of toxicity: a valuable feature in drug development. In addition to its selective index of ≥40 [10] and reports that it localised to erythrocytes in human trials [17,18], these characteristics could allay concerns about host-parasite specificity, which are often an issue when targeting eukaryotic parasites. Here, we set out to characterise the anti-parasitic mode of action for both quarfloxin and a related drug CX-5461 (brand-named Pidnarulex), which remains under investigation as an anticancer chemotherapy [25,26].

We report that quarfloxin and CX-5461 both interact with quadruplex-forming sequences of parasite DNA, and that they both cause some signatures of DNA damage. However, this may not be the sole mode of action, particularly for quarfloxin: CX-5461 is a more effective DNA damaging agent than quarfloxin, yet quarfloxin is more toxic to parasites than CX-5461. To gain more insight into this difference, we also explored the acute effects of both drugs on the trophozoite transcriptome and found distinct signatures of transcriptional disruption, with several PQS-containing genes affected. CX-5461 was heavily associated with disrupting replication and erythrocyte-invasion pathways, while quarfloxin disrupted fewer genes, including those in variantly-expressed sub telomeric gene families and those involved in the heatshock stress response. Overall, we find differing mechanisms of action for these compounds, although both function as DNA damaging agents in *Plasmodium*.

## Results

### Quarfloxin is more toxic to *Plasmodium* parasites than CX-5461

The EC_50_ of CX-5461 in *P. falciparum* parasites (3D7 strain) was measured and compared with the EC_50_ of quarfloxin, which we previously reported at 110 nM [10]. The EC_50_ of CX-5461, measured using a standard 48h Malaria SYBR-green Fluorescence (MSF) assay [27], was 440nM, meaning that *P. falciparum* is fourfold more sensitive to quarfloxin than CX-5461 (Figure 1A).

**Fig 1.**
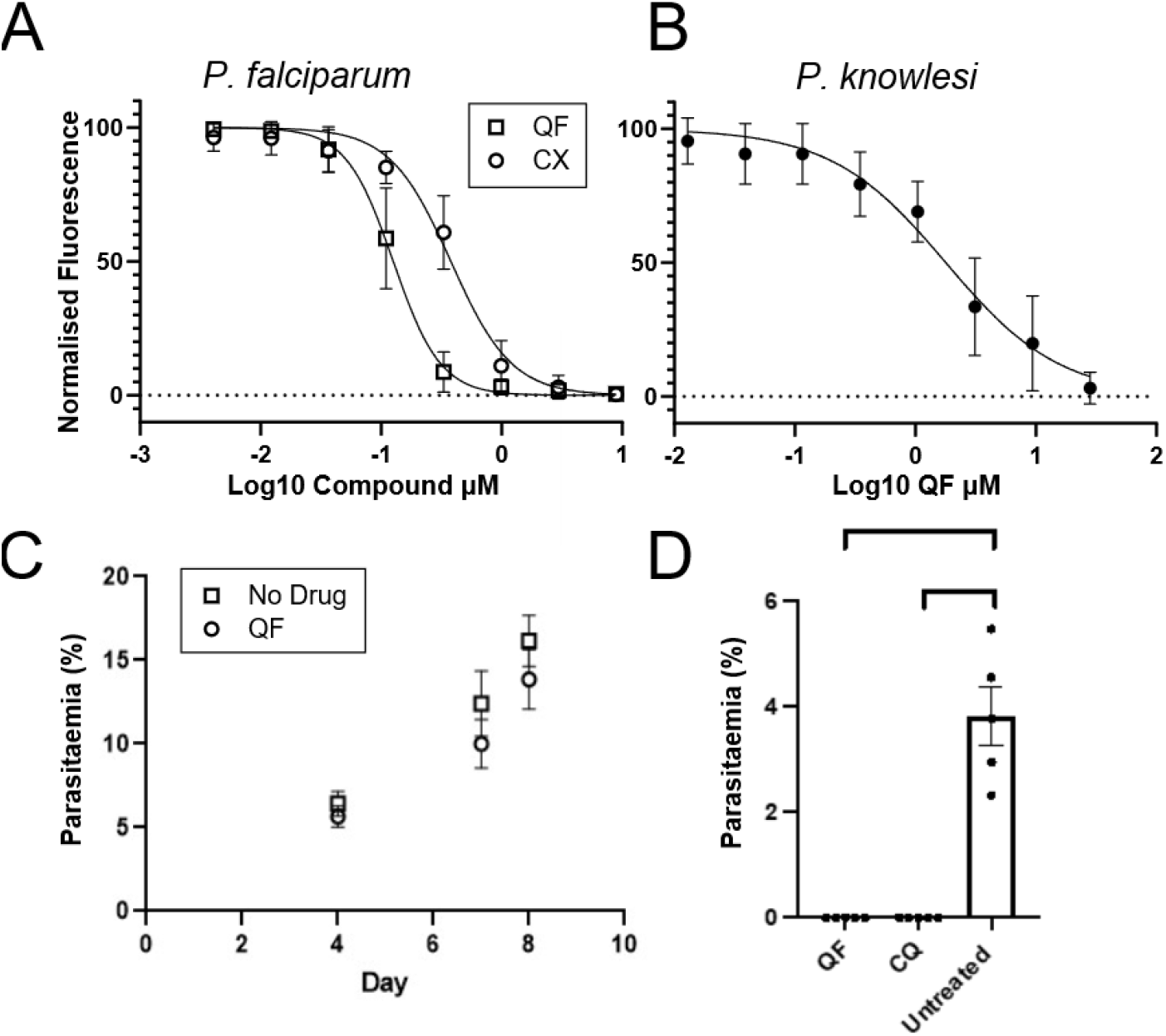
Quarfloxin and CX-5461 are toxic to several Plasmodium species. A) Dose response curves of quarfloxin and CX-5461 in P. falciparum and B) Dose response curve for quarfloxin in P. knowlesi. Curves were generated from MSF assays (3 biological repeats) and used to calculate EC_50_ values. P. falciparum was more sensitive to quarfloxin (EC_50_ =110 nM) than CX-5461 (EC_50_= 440 nM). P. knowlesi was less sensitive to both quarfloxin (EC_50_ =1.64 µM) and CX-5461 (which was limited by solubility, precluding the generation of a dose response curve at reasonable concentrations of the DMSO vehicle). C) Results of a preliminary in vivo Peter’s test using 0.6mg/kg quarfloxin administered IV to P. berghei-infected mice (n=5 per group). Despite a small reduction in parasite burden the mice did not clear infection over 8 days. D) Parasitaemia of mice (n=5 per group) 5 days post infection with 10^5^ P. berghei parasites and then daily treatment with 25 mg/kg quarfloxin administered IP. Chloroquine (CQ) is shown as a positive control for parasite clearance.

We also evaluated the relative toxicity of these drugs in the other readily cultured human malaria species, *P. knowlesi (*Figure 1B), since antimalarial drugs are particularly valuable if they work across species and *P. knowlesi* is often used as a model for the second major human parasite, *P. vivax*. The EC_50_ of quarfloxin in *P. knowlesi* was 1.64 µM, more than an order of magnitude higher than in *P. falciparum*. In fact, for many other antimalarials it has been reported that *P. knowlesi* is less sensitive *in vitro* than *P. falciparum* [28]. Nevertheless, this is still within the desirable range for developing new antimalarials (<2 µM) [29]. Satisfactory dose response curves could not be generated for CX-5461 in *P. knowlesi* without exceeding the solubility of the drug.

The more potent of the drugs, quarfloxin, was then evaluated in an *in vivo* murine model against the model parasite species *P. berghei.* An initial Peter’s test treated infected mice at estimated doses of 100x EC_50_ (0.6mg/kg, estimated crudely from the *in vitro* 48h EC_50_ measured on *P. falciparum*). The drug was administered through the intravenous route. Quarfloxin-dosed mice showed a slight decrease in parasite burden, as determined by tail-vein smears (Figure 1C), but there was no meaningful efficacy. The dose was then increased to 25 mg/kg with delivery by intraperitoneal injection (Figure 1D). Under these conditions, mice completely cleared all blood-stage parasitaemia at day 5 post-infection, as determined by examination of peripheral tail-vein smears. Previous patent documents for quarfloxin had suggested that the drug was well-tolerated in mice at this dosage [30], but equivalent levels administered in our experiments caused severe adverse effects, both via the intra-peritoneal route (necessitating euthanasia 5 days post infection) and via the intravenous route (causing severe inflammation at the injection site). Further experiments were therefore precluded, despite the promising effect of quarfloxin in killing *P. berghei*.

### *P. falciparum* cannot readily develop resistance to quarfloxin in *in vitro* culture

The risk of developing drug resistance is an important consideration for any novel antimalarial candidate. We therefore tried to develop quarfloxin-resistant parasites via long term *in vitro* culture of *P. falciparum* with continuous exposure to sub-lethal levels of quarfloxin, slowly increasing in concentration. After many months, clones could be generated with only modest (∼two-fold) reduced sensitivity to quarfloxin. MSF analysis did not show meaningful increases in EC_50_ values (Supplementary Figure 1), and resistance was not stable after short-term drug removal, suggesting that it was not genetically encoded. This failure to generate quarfloxin-resistant parasites meant that it was not possible to define a mechanism of action for the drug by identifying mutated resistance genes.

### Quarfloxin is distributed throughout *P. falciparum* parasite cells

The subcellular location of a drug can potentially help to reveal its mechanism of action. For example, chloroquine accumulates in the food vacuole [31] – the target organelle in which it inhibits haem detoxification. Therefore, the fluorescent nature of quarfloxin as a fluoroquinolone was exploited to assess its localisation within the parasite cell using live fluorescence microscopy. (Equivalent experiments with CX-5461 were precluded by its non-fluorescent nature.) Quarfloxin has an excitation maximum of approximately 350 nm and emission maximum of 550 nm. In conjunction with live-cell-compatible dyes emitting in the far-red range (DRAQ5, MitoTracker FR and Lysotracker FR), microscopy on live parasites could show any compartmentalisation of the compound to the nucleus, mitochondrion or food vacuole.

Quarfloxin preferentially localised within infected erythrocytes over empty erythrocytes and was confined within the parasite membrane in infected erythrocytes (Fig. 2). However, there was no specific intracellular concentration. Although quarfloxin signal could be seen in nuclei as anticipated (Figure 2A, red), signal was also detected around the mitochondria (Fig. 2A, magenta) and the food vacuole (Figure 2A, blue). Colocalisation analysis showed there was no colocalisation between quarfloxin and any organelle tested (Figure 2B). Signal was highly diffuse across the entire parasite cell (being excluded only from the inorganic haemozoin crystal in mature parasites) and it did not appear in foci.

**Fig 2.**
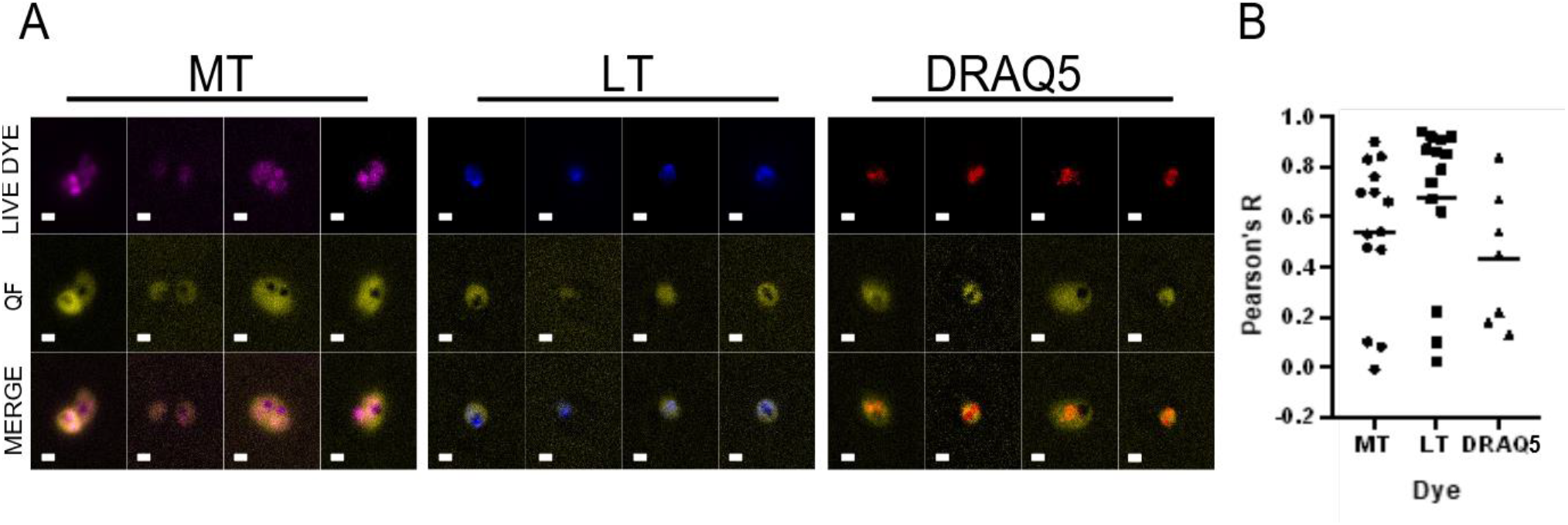
Localisation of quarfloxin within cells. A) Representative images of live parasites stained with quarfloxin (yellow) and MitoTracker (magenta, mitochondrial signal); LysoTracker (blue, food vacuole signal); or DRAQ5 (red, nuclear signal) live-cell-compatible dyes. Scale bar = 5 µm. B) Images were analysed for colocalisation of quarfloxin and organellar signal using ImageJ (n=∼10 cells). No obvious colocalisation within specific organelles was observed.

### CX-5461 does not target *P. falciparum* through topoisomerase inhibition

CX-5461, like quarfloxin, was originally thought to function as an RNA polymerase I inhibitor in human cells. Our earlier work already ruled out this mode of action for quarfloxin in *P. falciparum*, probably due to the non-canonical arrangement of rRNA genes in this parasite [10]. More recently, however, it was reported that the primary mechanism of action for CX-5461 in human cells is actually to cause DNA damage through topoisomerase II inhibition. Accordingly, in cancer cells, resistance to topoisomerase inhibitors tracks with resistance to CX-5461 [26]. This was equally possible as a mechanism for CX-5461 in *P. falciparum*: the parasite encodes well-conserved topoisomerases [32] and is sensitive to topoisomerase poisons such as etoposide [12].

We therefore evaluated the hypothesis that CX-5461 could affect *Plasmodium* topoisomerase activity, utilising an existing line with an altered capacity for DNA double-strand-break repair. This line was generated via a mutated *RAD54* gene [12]. The *P. falciparum RAD54* gene happens to encode an rG4 that reduces gene translation, so an rG4-mutant produces far more RAD54 protein than a wildtype parasite. Accordingly, the mutant line has reduced sensitivity to etoposide, presumably through more efficient DNA repair [12]. If CX-5461 acts similarly to etoposide, this mutant should also have reduced sensitivity to CX-5461. However, we found negligible change in the EC_50_ of CX-5461 in ‘RAD54-high’ parasites, in contrast to their differential etoposide sensitivity. The same parasites were actually somewhat less sensitive to quarfloxin, but only by about 1.5-fold compared to a 2.5-fold change in etoposide sensitivity (Supplementary Figure 2). Therefore, we could not generate consistent positive evidence for this mode of action in *Plasmodium*.

### DNA damage phenotypes are caused in *P. falciparum* by both quarfloxin and CX-5461

Despite the negative data on topoisomerase inhibition, it remained possible that quarfloxin and CX-5461 would cause DNA damage in *P. falciparum* by an alternative route, such as stabilizing G4s and hence stalling DNA replication forks. Experiments were therefore undertaken to explore the ability of both drugs to cause DNA damage in *P. falciparum*.

DNA strand breakage was assessed by terminal deoxynucleotidyl transferase dUTP nick end labelling (TUNEL) assay (Figure 3A-B). In this assay, broken or nicked DNA ends are labelled with a nucleoside that can then be imaged using fluorescence microscopy. Representative images (Figure 3A) showed some colocalisation of TUNEL and DAPI (DNA) stain in control cells, but this occurred in significantly more cells after treatment with CX-5461 (p = 0.0011), as well as after treatment with two positive controls, bleomycin (p = 0.0025) and DNase (p<0.0001), which cause single and double stranded DNA breaks. This indicated a causative relationship between all these drugs and DNA damage (Figure 3B). Quarfloxin-treated samples also had more co-stained nuclei than controls, but fewer than CX-5461-treated samples: TUNEL staining here was at a level similar to that in parasites treated with chloroquine, an antimalarial that does not damage DNA as its primary mechanism of action. Accordingly, neither chloroquine nor quarfloxin showed a statistically significant increase in TUNEL staining over the control.

**Fig 3.**
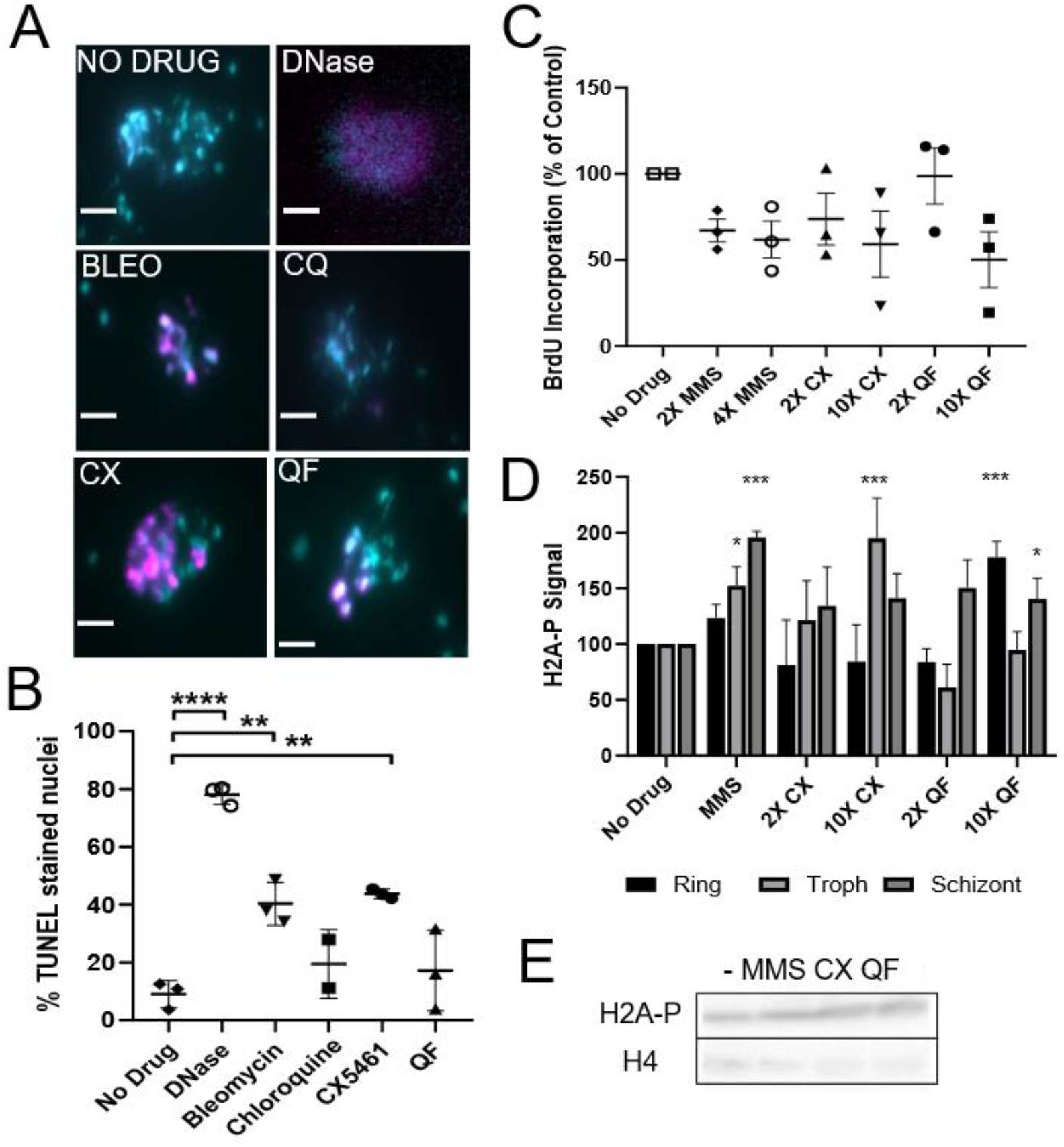
Quarfloxin and CX-5461 are associated with DNA damage. A) TUNEL staining of drug treated parasites. Mixed-stage cultures were incubated with drugs (2X EC_50_) for 4 h, stained with TUNEL kit (magenta), counterstained with DAPI (cyan) and imaged via light microscopy (Scale bar = 2 µm). DNase treatment and bleomycin were used as positive controls, and chloroquine (a known antimalarial that does not target DNA directly), as a negative control. B) Nuclei were counted and the percentages of nuclei co-stained with TUNEL and DAPI were calculated and plotted (3 replicates: n = 100 cells). Data were tested for significant differences against the untreated control via one way ANOVA with Dunnett’s posthoc. C) BrdU incorporation measured by ELISA. Parasites were incubated with high and low concentrations of drug (10X or 2X EC_50_) for 1 h and pulsed with BrdU for a following 1 h prior to ELISA assay. BrdU incorporation was measured as signal intensity as a percentage of signal in untreated controls. Three biological repeats were assessed for significant differences by one way ANOVA. D) H2A-P signal intensity as measured by densitometry on three biological repeats of western blots after parasites had been treated with 2X EC_50_ quarfloxin, CX-5461 or MMS for 2 h. 2-way ANOVA with Dunnett’s posthoc was performed. E) Representative western blot showing H2A-P signal; loading was controlled with histone H4 band density. All stars indicated significance at * P<0.05, ** P<0.01 ***P<0.001, P<0.0001.

Checkpoint responses to DNA damage are not well-characterised in *Plasmodium*, but in most eukaryotic cells, damage triggers a reduction in DNA replication – either directly by impeding replication forks, or via checkpoint responses that prevent S-phase entry (a G1-S checkpoint) or reduce the progress of an ongoing S-phase (an intra-S-phase checkpoint) [33]. Therefore, DNA replication was measured after treatment with quarfloxin and CX-5461, using nascent DNA labelling (Figure 3C). Parasites made to express a viral thymidine kinase enzyme can scavenge pyrimidine nucleosides during replication, and thus nucleoside labelling techniques, which are otherwise unavailable in *Plasmodium*, can be leveraged to quantify nascent DNA replication [34]. Here, the nucleoside analogue bromodeoxyuridine (BrdU) was used to label synchronous trophozoite parasites (this being the cell-cycle stage in which S-phase occurs) for 1h directly after a 1h pulse of drug. We used high (10X EC_50_) or low (2X EC_50_) concentrations of quarfloxin, CX-5461, or the control DNA-damaging agent methylmethanesuphonate (MMS). MMS is well-characterised in model cells to alkylate DNA and thus acutely arrest ongoing replication forks [35,36]. Nascent DNA synthesis was measured by BrdU ELISA [34]. DNA replication was acutely reduced after DNA damage with MMS and also quarfloxin and CX-5461. This response was more pronounced in CX-5461, where both low and high levels of drug reduced DNA replication; quarfloxin required higher levels (10X EC_50_) to achieve similar disruption of replication.

The DNA damage response in many eukaryotic cells is characterised by phosphorylation of the H2AX histone [37]. *P. falciparum* lacks H2AX but was recently reported to make an analogous DNA damage response by phosphorylation of H2A [38]. This can be observed using an antibody to phosphorylated human H2AX, which is cross-reactive against phosphorylated *P. falciparum* H2A (Figure 3D, E). Western blots were performed on lysates collected from synchronised ring, trophozoite and schizont parasite cultures after 2h drug treatments, to assess if there were stage-specific differences in DNA damage signalling via H2A-P. The signal appeared to be both stage dependent and dose dependent. MMS and CX-5461 both elicited phosphorylation primarily in trophozoites and schizonts, consistent with DNA damage occurring in a replication-linked fashion. Quarfloxin gave less consistent results but at higher levels it elicited the signal in rings and schizonts but not, unexpectedly, in trophozoites. In all cases, the increase in H2A-P signal was modest (under 2-fold) and also transient, peaking within ∼2h (data not shown).

As both quarfloxin and CX-5461 generated DNA damage hallmarks, we tested if they may be acting in the same pathway by isobologram analysis [39,40] (Supplementary Figure 3). We found that the drugs did not synergise with one another, with drug combinations falling on or above the additive line, providing weak evidence that they may compete antagonistically for the same targets.

### QF and CX-5461 bind *in vitro* to *P. falciparum* G-quadruplexes

The data in figure 3 raised the question of exactly how DNA damage is caused by quarfloxin and CX-5461. Supplementary figure 2 has already showed that it is unlikely to be primarily via topoisomerase inhibition. However, since these drugs are designed to bind to G4s, they could hypothetically stabilise such structures, impede DNA replication forks and thus cause fork stalling and DNA breakage [41–43]. There is no direct evidence for this as yet in *P. falciparum*.

Oligonucleotides encoding *P. falciparum* G4 sequences were used to assess the G4 binding capacity of quarfloxin and CX-5461 *in vitro* (Figure 4). Oligos were folded *in vitro* and their correct folding was confirmed by incubation with Thioflavin T (ThT), a small molecule that fluoresces only when bound to G4 structures [44]. Two different G4 structures were selected, alongside controls with a scrambled sequence, or a sequence of A/T-only, neither of which bound to ThT (Figure 4A, B). A titration of either quarfloxin or CX-5461 was added to the ThT-bound oligos to test whether these drugs could bind to G4s and outcompete ThT, resulting in a ‘quenched’ signal. Both quarfloxin (Figure 4A) and CX-5461 (Figure 4B) could compete with ThT, even at lower-than-1:1 molar ratios of drug:ThT.

**Fig 4.**
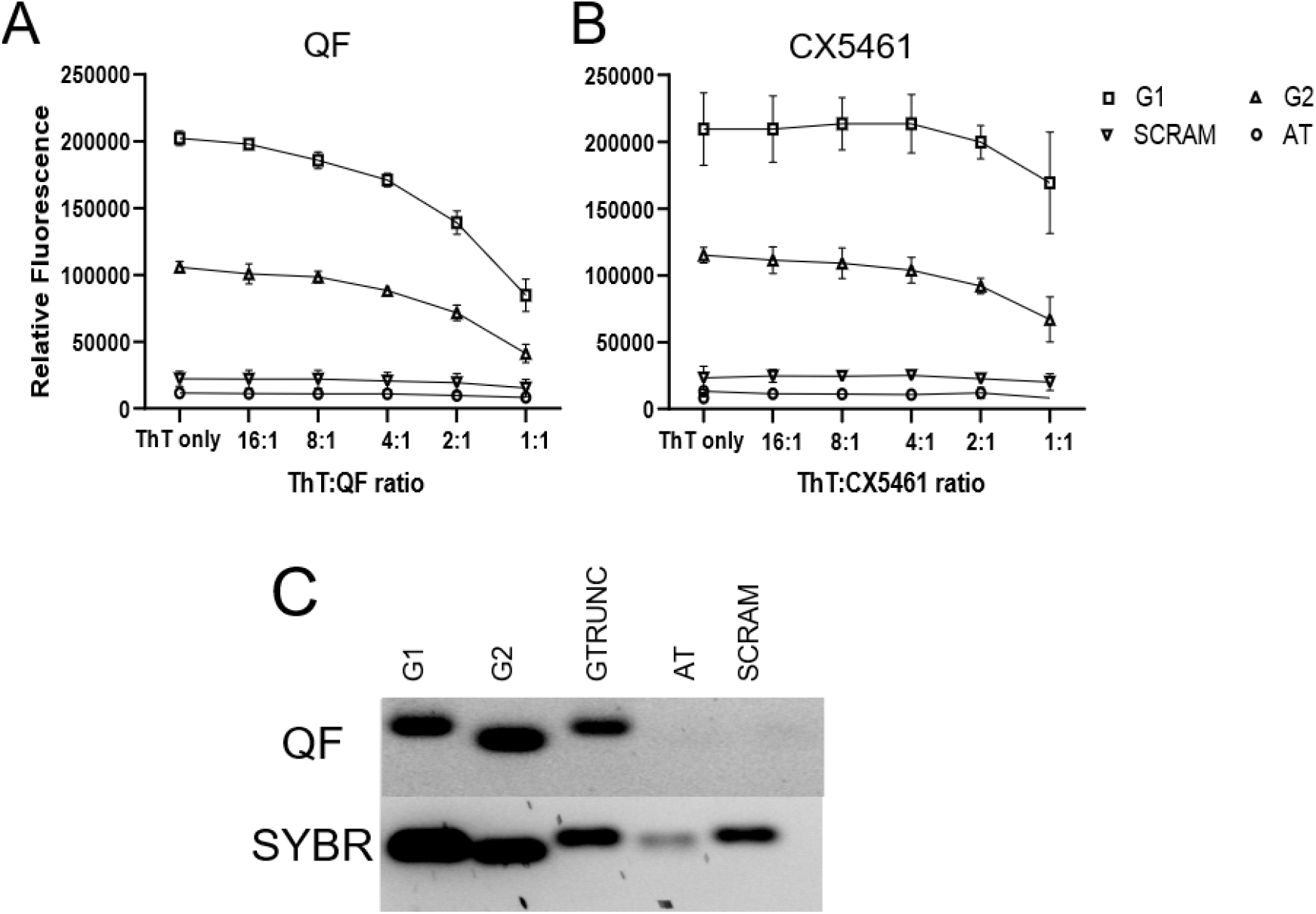
Quarfloxin and CX-5461 can bind to G4 sequences found in P. falciparum. Oligo sequences were allowed to fold into G4s by annealing in the presence of monovalent cations and then incubated in the presence of 40 µM ThT and a dilution gradient of either A) quarfloxin or B) CX-5461. Oligos were incubated for 30 min before reading the ThT fluorescence as relative fluorescence units (RFU). Average RFUs were calculated from three blank-corrected biological repeats. Oligo sequences were G4-encoding (G1, G2), scrambled (scram) or A/T-only (AT). C) Visualisation of quarfloxin binding by agarose gel electrophoresis. 10 µM oligos incubated with 100 nM quarfloxin were imaged under UV (for quarfloxin) and blue light (for SYBR green).

Due to its fluorescent property, the binding of quarfloxin to G4 folded oligos could also be visualised on an agarose gel (Figure 4C). Incubation of 10 µM folded oligo with 100 nM quarfloxin showed preferential binding to the G4 folded oligos. A truncated oligo bound more weakly (possibly by forming inter-molecular G4s) and scrambled or A/T-only oligos did not bind at all.

### QF and CX-5461 do not detectably stabilize *P. falciparum* G-quadruplexes *in cellulo*

The data in figure 4 showed that both drugs can bind to parasite derived G4s *in vitro,* but not that they actually do so *in vivo.* The BG4 antibody [45] has been used to visualise G4s in fixed cells in immunofluorescence assays [46–48], including in *P. falciparum* [10], and more recently in flow cytometry [49]. Cultured parasites were incubated with quarfloxin or CX-5461 and analysed by immunofluorescence (Figure 5A). BG4 signal was present primarily in nuclei, as previously reported [10], suggesting that the signal is more representative of DNA G4s than of RNA G4s – indeed, the great majority of PQSs in the *P. falciparum* genome are found in telomere repeats. Quarfloxin and CX-5461 did not significantly affect the BG4 signal intensity (Figure 5B). Parasites were also assessed by flow cytometry for a more quantitative measurement (Figure 5C), but again no significant difference was observed (Figure 5D).

**Fig 5.**
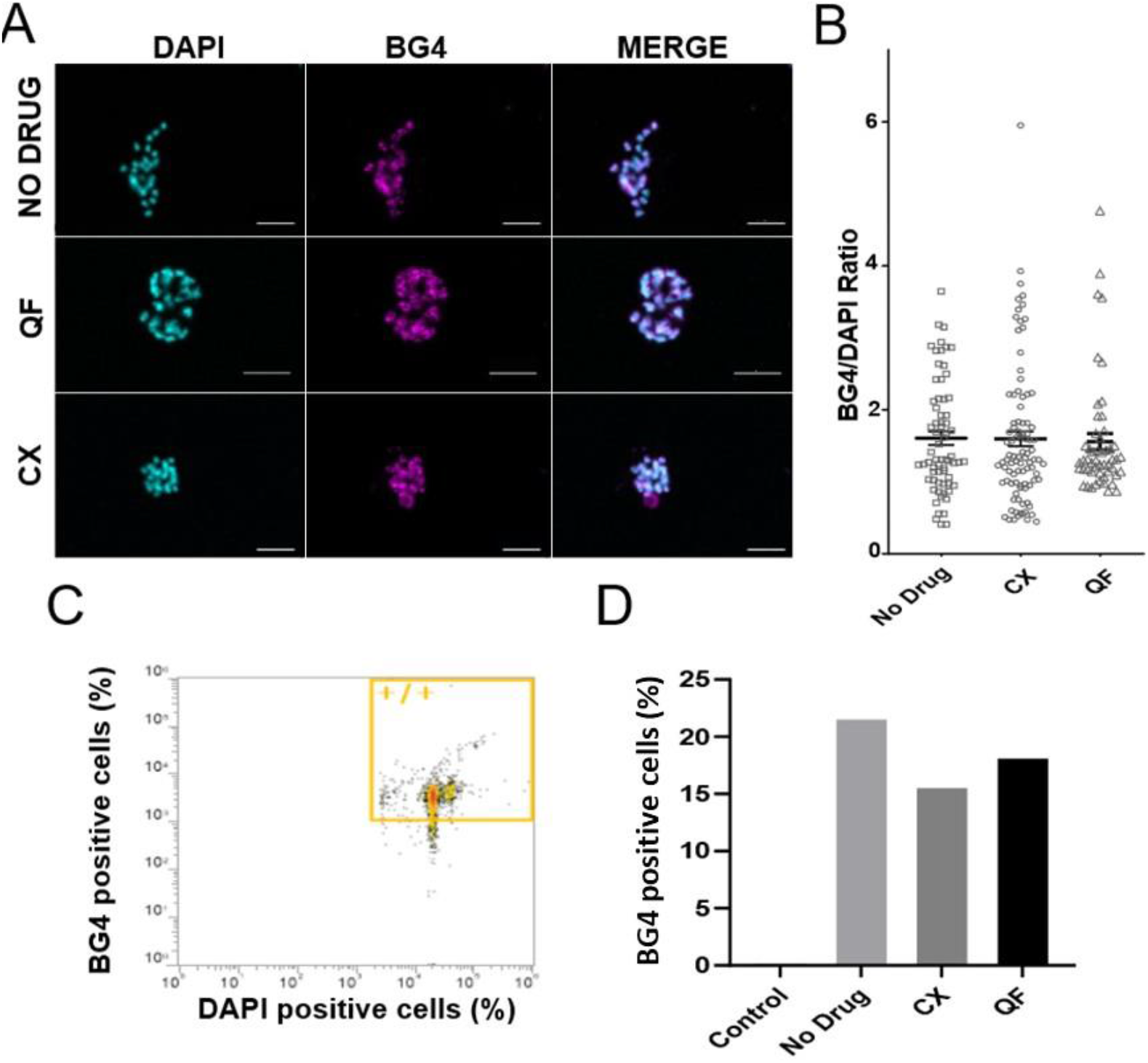
Quarfloxin and CX-5461 do not alter the total G4 signal detected in cellulo. A) P. falciparum cultures were incubated in no drug, 2X EC_50_ quarfloxin or CX-5461 for 4 h prior to analysis by immunofluorescence. Cells were stained with DAPI (cyan) and BG4 (magenta) and imaged by confocal microscopy. Scale bar = 5 µm. B) Signal intensity per cell was measured in ImageJ and represented as BG4 intensity relative to DAPI signal (n = ∼50 cells). C) Flow cytometry analysis was performed, recording 20,000 events, gated for singlets, and then for +/+ BG4/DAPI signal. Example data are shown from cells treated with CX-5461. D) The percentage of BG4-positive cells in a population of 20,000 is shown after treatment with quarfloxin, CX-5461 or no drug.

We previously generated mutants in two *P. falciparum* RecQ helicases, termed *Pf*BLM and *Pf*WRN. These helicases play key roles in unwinding DNA G4s, and the mutant lines showed increased sensitivity to the G4-stabilising agent TMPyP4, as expected if they were impaired in their ability to unwind drug-stabilized G4s [50]. The same parasite lines were tested here, but they were only marginally sensitised to quarfloxin or CX-5461 (Supplementary Figure 4): a small shift in EC_50_ was observed, but this was not significant. Collectively, although quarfloxin and CX-5461 could clearly bind to G4 structures *in vitro*, the evidence was inconclusive for any broad G4-stabilising effect *in cellulo*.

### QF and CX-5461 have acute effects on the *P. falciparum* transcriptome

Since it was not possible, at a whole-cell level, to detect any bulk effects of quarfloxin or CX-5461 on *Plasmodium* G4s, we sought to improve our resolution and detect any gene-by-gene effects by exploring the potential of these compounds to affect the transcriptome. Transcriptome-wide RNA-Seq, optimised for *P. falciparum* [51], was performed on trophozoite parasites after treatment with no drug, quarfloxin or CX-5461 at 2X EC50 for 4h, between 30-32h and 34-36h post invasion (h.p.i). At this stage, the parasites should be performing S-phase. A moderate level and short period of treatment were chosen to capture only the most acute drug-specific effects, rather than any general changes associated with parasite death.

Both drugs caused significant transcriptional deregulation (Figure 6A-B) although greater numbers of genes were differentially expressed (with a fold-change > 2, FDR < 0.05) after CX-5461 (218 genes, Figure 6B) than after quarfloxin (51 genes, Figure 6A) (Figure 6C, Supplementary Table 1-2). Quarfloxin elicited very little upregulation, but downregulation of multiple genes encoding ncRNAs, including some ribosomal RNA fragments, and *PIR* genes, a family of sub-telomeric genes associated with virulence. Downregulated genes in CX-5461 included several ncRNAs (again including some ribosomal RNAs, but again without a strong, comprehensive effect suggestive of RNA polymerase I inhibition), rhoptry and apical-complex associated genes, and genes for merozoite surface proteins (MSPs). Upregulated genes included *PIR*s (in direct contrast to the downregulatory effect of quarfloxin), and heatshock associated genes (Supplementary Table 2). Overall, genes were more likely to be downregulated than upregulated by both compounds. Around 34% of genes modulated by CX-5461 had unknown functions, and ∼25% for quarfloxin, which is not unusual in the *Plasmodium* genome.

**Figure 6.**
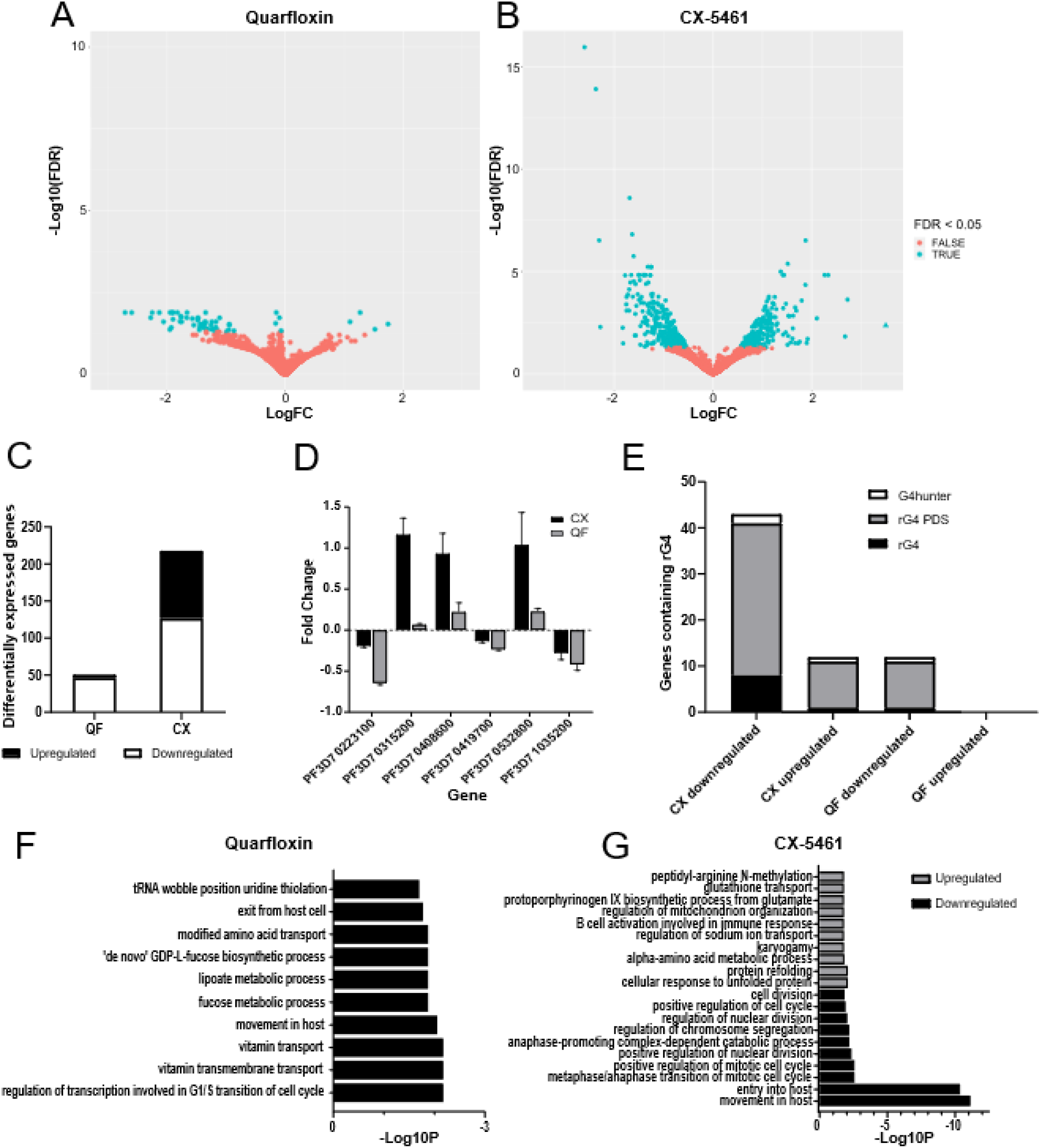
Quarfloxin and CX-5461 dysregulate the parasite transcriptome. A,B) Volcano plots of differentially expressed (DE) genes (adj. P < 0.05, log2FC >1) for A) quarfloxin and B) CX-5461 compared to untreated control. C) Bar chart of total numbers of up- and down-regulated DE genes after each drug treatment. D) Validation of RNA-Seq by qRT-PCR on six selected genes after parasites received the same treatment with quarfloxin or CX-5461. The qRT-PCR experiments were repeated in biological triplicate and expression of each gene was calculated versus the average of two housekeeping genes. E) Bar chart of the number of genes dysregulated by either drug that also contained rG4s (generated by comparison with a published rG4-seq dataset of rG4s detected in the presence or absence of pyridostatin) or DNA G4s (by comparison with published predictions from G4Hunter). F, G) GO enrichment analysis was performed. Top 10 biological processes amongst DE genes are plotted for F) quarfloxin downregulated DE genes and G) CX-5461 up-and down-regulated DE genes.

To further validate the RNA-Seq data, qRT-PCR was performed on a small set of 6 modulated genes, selected because they had large fold changes (so would be easier to detect by qRT-PCR) or had low P values (potentially more biologically relevant). All but one of these genes (PF3D7_0315200 for quarfloxin) displayed a positive or negative fold change concordant with the RNA-Seq data (Figure 6D).

We assessed the G4 content of deregulated genes using several published datasets of genes predicted to encode G4s [9, 12]. Concerning RNA G4s, 10 quarfloxin-modulated genes and 43 CX-5461-modulated genes encoded putative rG4s (Figure 6E, Supplementary Table 3) [12]. The rG4-encoding genes were largely downregulated. They accounted for 19.6% of all quarfloxin-modulated genes and 19.7% of CX-5461-modulated genes; however, these proportions are similar to the proportion of all *P. falciparum* genes that encode putative rG4s when stabilised by the drug pyridostatin (22%), so there was no clear enrichment of rG4-encoding genes. By comparison, only a few drug-dysregulated genes containing PQSs in their DNA sequence, as predicted by G4Hunter [9] (Supplementary Table 3).

In addition to directly regulating G4-encoding genes, it was possible that quarfloxin and CX-5461 would ‘indirectly’ affect particular transcriptional pathways, as the parasites made an acute response to drug stress. Gene ontology (GO) enrichment analysis was performed using ReviGO [52] to detect any enriched gene activities amongst the dysregulated genes (Figure 6F-G, Supplementary Figure 5). Quarfloxin caused very little upregulation of genes and so analysis was performed on downregulated genes only. Quarfloxin treatment was associated with downregulation of processes related to vitamin transport, mitotic cell cycle transcription and movement in the host (Figure 6F). Enriched molecular functions included peroxidase and sulfotransferase activity, DNA exonuclease activity and NAD dependent histone deacetylation (Supplementary Figure 5A). These pathways, however, were generally represented by only small numbers of genes. CX-5461 treatment was characterised by downregulation in biological processes related to cell cycle regulation, nuclear division, mitotic processes, and parasite entry into the host (Figure 6G). Further enrichment was observed in GO terms related to cell surface and NAD^+^ binding, peroxidase activity, and transmembrane transport (Supplementary Figure 5B). Upregulated genes from CX-5461 treatment were enriched for GO terms associated with protein refolding, misfolded protein rebinding, programmed cell death and ion transport regulation (Figure 6G, Supplementary Figure 5C).

We performed further GO enrichment analysis on a larger group of genes showing a less-stringent fold change of 1.5 rather than 2 (Supplementary Figure 6). This did not have any major effect on the GO terms for genes downregulated by quarfloxin or CX-5461, but it did seem to increase diversity of terms in genes that were upregulated following CX-5461 treatment. There was a notable increase in terms pertaining to ribosomal biogenesis and RNA transcription initiation compared to more stringent 2-fold expression analysis.

## Discussion

This work explored the potential of two G4-binding drugs – previously developed and tested as anticancer agents in humans – for repurposing as antimalarials. It also sought to establish a molecular mechanism of action for these drugs in malaria parasites. We found that although quarfloxin and CX-5461 may be unsuitable for repurposing in their current form, they do provide insights into the potential drugability of G4s in *Plasmodium*, which is a major human pathogen and also an unusual, early diverging eukaryote with a guanine-poor genome.

We previously reported that quarfloxin had fast and potent parasite-killing activity *in vitro* [10]. Here we showed that it is also active *in vivo* in a mouse model. This represents an important step in drug development, but high quarfloxin treatments severely debilitated the mouse host as well as the parasite. Although the original patent reported tolerance in mice, and although the selective index of quarfloxin on *P. falciparum* versus cultured mammalian cells is theoretically large, it evidently has off-target toxicity in mice. Our data, for the first time, prove the principle that G4-targetting is a viable antimalarial strategy *in vivo*, but drugs with lower toxicity are clearly needed. When we tested the more recently developed CX-5461, which is still undergoing human cancer trials and is adequately tolerated in humans [25], we found this to be 4-fold less potent against *Plasmodium* than quarfloxin.

Adding confidence to G4-targetting as a general antimalarial strategy, quarfloxin killed the zoonotic *Plasmodium* species *P. knowlesi* as well as *P. falciparum*. Its apparently lower potency in *P. knowlesi* may be a technical artefact, because *P. knowlesi* is cultured in high concentrations of horse serum that reduce the effective concentration of many drugs [28,53]. Alternatively, it may highlight a genuine difference in the biology of the two species: *P. knowlesi* has a G/C-balanced genome and hence a higher density of PQSs than *P. falciparum* [9,54]. Perhaps, counterintuitively, this renders parasites less sensitive, rather than more sensitive, to G4 binding. To pursue this issue, a whole range of G4-binding drugs would need to be tested in *P. knowlesi*.

In *P. falciparum* both compounds had steep dose response curves: quarfloxin in particular had a steep curve characterised by high *m* values, indicating greater potency in small increments above the EC_50_. This hallmark was present across species for both compounds, although to a greater extent in quarfloxin. Published work has shown that two compounds with the same EC_50_ value but differing *m* values can have different inhibitory effects, with a steeper slope associated with better inhibitory effects at clinical levels [55].

Another important issue for drug development is the selection of resistance, which historically has sometimes occurred within months of clinical use of a new antimalarial drug [56,57]. Single enzyme targets can be most problematic in this regard, whereas ‘irresistible’ drugs are thought to have complex/multifactorial modes of action. Resistance can potentially be recapitulated in culture, which is useful for identifying mutated gene targets by sequencing resistant parasites [58]. Quarfloxin, however, proved ‘irresistible’ in culture, suggesting that it may have a complex mode of action and also precluding any identification of drug target(s) from resistant parasites. Similarly, the failure of quarfloxin to accumulate in any particular parasite organelle added no clues as to its site of action.

Since all these agnostic approaches failed to elucidate how quarfloxin kills *Plasmodium*, we turned to hypothesis-driven experiments, based around the induction of DNA damage. DNA damage or replication stress may be a particularly effective antimalarial strategy because *Plasmodium* replicates fast and extensively during schizogony, and because it lacks the nonhomologous end-joining pathway for DNA repair and is entirely reliant on homologous recombination [59]. Indeed, antifolate drugs that inhibit DNA replication are very effective antimalarials (albeit subject to resistance in the field) [59].

Quarfloxin and CX-5461 could theoretically cause DNA damage either via topoisomerase II (as evidenced for CX-5461 in human cells [26]), or via G4 stabilisation and replication fork stalling (evidenced for several G4-binding drugs in model systems [41,60,61]). We found little consistent evidence for the former: although our experiments did not categorically rule it out, they at least showed that these drugs do not act exactly like etoposide. We did, however, find evidence for the latter: both drugs induced some level of DNA breakage; both induced acute slowing of DNA replication that is highly consistent with fork stalling; and both induced phosphorylation of a damage-responsive histone. Responses occurred fast – within 1-2h of drug treatment – making it likely that they were direct drug effects, not merely side-effects of cell death. All responses were stronger with CX-5461 than with quarfloxin (despite CX-5461 having no demonstrable effect through topoisomerase II). Hence, we concluded that CX-5461 may kill parasites primarily through DNA damage, but that quarfloxin could be more complex, since it is more toxic than CX-5461, yet less DNA-damaging.

Pursuing this mechanism, we showed that quarfloxin and CX-5461 can bind *in vitro* to *Plasmodium* G4 sequences, displacing ThT (which is an end-stacking molecule – so quarfloxin and CX-5461 probably also end-stack on G4s). *In cellulo*, however, we detected no global increase of G4s in drug-treated parasites. In fact, the antibody-detectable G4 signal in *P. falciparum* is probably heavily dominated by G-rich telomeres, and these may not be the relevant target of these drugs, since we previously showed that sub-lethal quarfloxin has no effect on telomere maintenance [10].

Finally, we used RNA-Seq to identify gene-specific targets of the two drugs, hypothesising that a) genes encoding G4s that are directly bound by the drugs could be repressed, and b) genes in any stress pathways acutely affected by the drugs would hint at the type of stress induced. We used a short, relatively low-level drug treatment to avoid non-specific effects, and accordingly found relatively few deregulated genes. (In a single precedent for this type of experiment, pyridostatin affected ten times as many genes, but parasites were exposed for an entire 48h cycle [8]. In a more comparable experiment, etoposide treatment for 6h deregulated only 225 genes [62].)

Among the genes affected by quarfloxin or CX-5461, only a minority encoded G4s – primarily RNA rather than DNA G4s. Our protocol would only detect transcripts that are turned over within 4h, but with this caveat, neither drug seems to kill parasites mainly through binding to a large cohort of essential G4-encoding genes and impeding their transcription or inducing transcript degradation. Nevertheless, the affected G4-encoding genes were frequently affected by *both* drugs, supporting G4 binding as one mode of action for these drugs, and suggesting that where they do bind G4s, they bind to similar structures. Isobologram analysis of the interaction between quarfloxin and CX-5461 also revealed weak antagonism, supporting an element of competition between the two drugs. The particularly low representation of PQSs at the DNA level, as predicted by G4Hunter even at a low stringency threshold, probably reflects the fact that *P. falciparum* encodes relatively few canonical G4s, whereas it encodes many more non-canonical motifs that can fold in RNA (as detected by rG4-Seq) but probably not in DNA.

RNA-Seq primarily detected early downstream effects of drug treatment: for CX-5461, a downregulation of replication and invasion pathways, consistent with the parasites detecting DNA damage and arresting their cell cycle and the maturation of daughter cells until damage can be repaired. This was accompanied by upregulation of heatshock pathways – a striking parallel with the response to the frontline antimalarial artemisinin. Artemisinin causes broad-spectrum damage to both proteins and DNA [62] and elicits a transcriptional response similar to the heatshock/misfolded protein response [63]. For quarfloxin, the same pathways were not affected: overall we saw fewer deregulated genes and they did not point clearly to a route to parasite killing. We then relaxed the fold-change threshold to 1.5 and repeated the GO enrichment analysis to capture any pathways affected at a lower levels, but the enriched pathways did not change greatly. Interestingly, a secondary enrichment appeared in the CX-5461-upregulated genes, pertaining to RNA transcription initiation and associated processes.

Overall, even this detailed analysis at the cellular, genomic and transcriptomic levels did not reveal a single clear mode of action for quarfloxin in *P. falciparum.* We found some evidence for DNA damage, but not primarily through topoisomerase inhibition, widespread G4-stabilization or telomere dysregulation. The high antimalarial potency of quarfloxin, which was seen both *ex-vivo* in cultured parasites and *in vivo* in mouse models, is probably due to polypharmacology: a positive feature for an antimalarial drug, but one that nevertheless remains mysterious.

## Methods

### Drug preparation and storage

Compounds were solubilised and stored in the dark at −20°C: quarfloxin at 1mM in ultrapure water, CX-5461 at 1.5 mM in DMSO.

### Parasite culture and lines

*P. falciparum* 3D7 parasites were cultured *in vitro* in customised RPMI 1640 medium (2.3 g/L sodium bicarbonate, 4 g/L dextrose, 5.957 g/L HEPES, 0.05 g/L hypoxanthine, 5 g/L Albumax II, Invitrogen) supplemented with 200 mM L-glutamine, 25 µg gentamicin and 2.5% human serum. Parasites were maintained at 4% haematocrit in human O^+^ erythrocytes at 37°C under specific gas mixture (3% O_2_, 5% CO_2_, 92% N_2_). *P. knowlesi* were maintained similarly, supplemented with 10% horse serum instead of human serum and 22.2 mM glucose, and maintained at 2% haematocrit. The Rad54 WT and Mut lines [12], *Pf*BLM and *Pf*WRN knockout/knockdown lines [50] and the strain expressing thymidine kinase [34] are previously described. For long term quarfloxin tolerance studies, 3D7 parasites were constantly cultured in the presence of sublethal doses of quarfloxin as determined by previous EC_50_ calculations [10]. Tolerance was checked at intervals of several months by new MSF assays. Cultures that displayed promising quarfloxin tolerance were subject to clonal dilutions and then repeated MSF assays.

### *P. berghei* experiments

All procedures were performed in accordance with the UK Animals (Scientific Procedures) Act (PP8697814) and approved by the University of Cambridge AWERB. The Office of Laboratory Animal Welfare Assurance for the University of Cambridge covers all Public Health Service supported activities involving live vertebrates in the US (no. A5634-01). This study was carried out in compliance with the ARRIVE guidelines (https://arriveguidelines.org/). Naïve CD1 mice (Charles River) (n = 5 per treatment) were infected by IP injection of 1 × 10^5^ *P. berghei* ANKA 2.34 parasites obtained from the blood of a donor mouse. Quarfloxin was dissolved in dH_2_O, and chloroquine (positive control) was dissolved in 7% Tween 80/3% Ethanol in dH_2_O and dosed either intraperitoneally or intravenously to infected mice to result in a 25 mg/kg exposure for both drugs. Five additional infected mice received a vehicle-only dose as a negative control. Infected mice then received identical doses of drugs/vehicle 0, 24, 48 and 72 h after infection and, commencing at 96 h post-infection, were smeared daily to monitor parasitaemia. Mice reached their humane endpoint immediately upon showing outward pathological indicators of malarial infection and a smear positive for malaria parasites. (They would be considered cured if this was not reached by day 30 post-infection). Mice dosed IP with 25 mg/kg quarfloxin demonstrated no sign of blood stage parasitaemia 96 h post-infection, but reached severity limits and were humanely euthanised at this point.

### Parasite synchronisation

Synchronised parasites were obtained by synchronisation 40 h apart with 5% sorbitol (w/v) to collect ring stages, or 65% v/v percoll gradient (GE Healthcare) to collect schizonts. Tightly synchronised parasites (2 h window) were created by incubating late schizont stages with 1.5 µM PKG inhibitor 4-[7-[(dimethylamino)methyl]-2-(4-fluorphenyl)imidazo[1,2-α]pyridine-3-yl]pyrimidin-2-amine (compound 2) [64] for 2 h before washing off and reinvading for 2 h in 25% haematocrit red blood cells as described previously [65].

### MSF Assay

Malaria SYBR green I based fluorescence (MSF) assay was performed as previously described [27]. 96 well black walled plates (Greiner) were predosed with a 3X serial dilution of relevant drug and trophozoite cultures were added in triplicate to a final 0.5% parasitaemia, 2% haematocrit. Each plate contained triplicate wells without drug as control, as well as triplicate wells of 2% haematocrit suspension without parasites for background correction. Outer wells contained culture media to prevent edge evaporation. Plates were incubated at 37°C for 48 h. Following incubation, samples were mixed 1:1 with 2X MSF lysis buffer (20 mM Tris, pH 7.5, 5.5 mM EDTA, 0.008% saponin, 0.8% Triton-x-100) containing 0.2 µL/mL SYBR green I (Sigma) and transferred to a black walled plate. Following 1 h incubation at room temperature in the dark, fluorescence was detected using a POLAR Star OMEGA (BMG) microplate reader (ex 490 nm; em 510-570nm). Normalised dose response curves and EC_50_ values were generated from blank-corrected means from three biological repeats using GraphPad Prism (v8) software (GraphPad Software Inc., San Diego, CA).

### Detection of quarfloxin in live-cell microscopy

Live (unfixed) *P. falciparum* parasites were incubated with 1 µM quarfloxin and either 50 nM DRAQ5 Abcam) or 100 nM Lysotracker DeepRed (ThermoFisher) or MitoTracker Deep Red FM (ThermoFisher) for 30 minutes in the dark at 37*°*C, washed, diluted 100-fold in PBS and mounted onto SecureSeal hybridisation chambers (Grace Biol-Labs). Parasites were imaged using a Nikon Eclipse Ti-E inverted microscope, temperature-controlled at 37°C, with a Nikon Plan Apo VC 60x/1.40 Oil objective and a CMOS camera (model GS3-U3-23S6M-C, Point Grey Research/FLIR Integrated Imaging Solutions (Machine Vision), Ri Inc., Canada). Quarfloxin was excited via Luxeon UV LHUV-0380-0200, peak at 380nm, using the following filters: Semrock FF01-390/40 excitation filter, Semrock FF520-Di02 dichroic, Semrock FF01-542/27 emission filter. The companion dyes, DRAQ5, Lysotracker DeepRed and MitoTracker Deep Red, were imaged with standard filter sets for these far-red dyes. Colocalisation analysis was performed in ImageJ.

### TUNEL assay

Parasites were incubated with 2X EC_50_ drug for 4 h and thin blood smears were created on glass slides. Samples were left to air dry, and then processed using *in situ* Cell Death Detection Kit TMR red (Roche) according to kit conditions and counterstained with 5µg/mL DAPI prior to mounting with Prolong Diamond antifade mountant (Invitrogen). Samples were imaged using a Nikon Microphot SA microscope using blue (DAPI) and red (TMR-red) filters at 60x magnification. Nuclei were counted and percentages of TUNEL stained nuclei were calculated. 80-100 nuclei were counted for each treatment and three biological repeats were performed.

### Western blot

Synchronous cultures at either ring, trophozoite or schizont stages were exposed to 2X EC_50_ drug for 2 h and parasites were saponin-released, lysed in RIPA buffer (150 mM NaCl, 50 mM Tris HCl, 1% v/v NP-40, 1 mM EDTA, 0.5% w/v sodium deoxycholate, 0.1% v/v SDS, 0.01% w/v sodium azide, pH 7.4), mixed with 4X LDS buffer (BioRad) and 5% v/v 2-mercaptoethanol and boiled for 5 mins at 95°C. Samples were electrophoresed on a 4-12% denaturing polyacrylamide gel (BioRad) in TGS buffer at 10 V/cm. The gel was transferred to a nitrocellulose membrane (Amersham) for 60 mins at 60V in chilled Towbin buffer. Following 1 h blocking with 3% BSA (w/v) 0.1% TBS-Tween20, the blot was probed overnight with either 1:1000 anti-phospho-histone-H2A.X Ser139 (#9718, CellSignalling) or anti H4 (ab10158, Abcam), rinsed, then probed with 1:2000 goat anti rabbit IgG-HRP (Abcam). Blots were developed using Clarity Western ECL substrate (Bio-Rad) and imaged using a GelDoc imager (Azure Biosystems). Following background correction and loading normalisation, densitometry analysis was performed in ImageJ [66] and the percentage increase in H2A-P signal was calculated relative to untreated samples.

### ELISA

Early trophozoite parasites were pulsed with high (10X EC_50_) and low (2X EC_50_) drug doses for 1 h and chased with 100 µM BrdU for 1 h. Parasites were saponin released from erythrocytes, washed and resuspended in PBS and then aliquoted in triplicate into clear flat bottom 96 well plates and left to dry at 37°C. A black walled plate was aliquoted with the same volume of parasites, mixed with an equal volume of 2X MSF lysis buffer containing 0.2 µl/mL SYBR green I and left to lyse for 1 h in the dark prior to reading on a platereader as per MSF assay protocol. This DNA measurement was then used to normalise the ELISA results for genomes loaded per well. After drying, parasites for ELISA were fixed for 15 mins in 4% PFA and then 1:1 methanol: acetone for 2 mins. Wells were rinsed three times with PBS for 5 mins before blocking for 1 h in 1% BSA (w/v) PBS. BrdU was detected with 1:500 mouse anti BrdU (MoBU sc-51514, Santa Cruz) and 1U nuclease in 1% BSA. The plate was sealed and incubated for 1 h at 37°C. Following 3 x 5 min rinses with PBS, wells were incubated with 1:5000 goat anti-mouse HRP (Dako) and incubated for 1 h in the dark. Following 3 x 5 min PBS rinses, TMB peroxidase (BioRad) was added and the plate was imaged on a POLAR Star Omega at 620nm following 5 and 10 mins development.

### Thioflavin-T quadruplex-binding assays

ThT G4 binding assays were performed as described previously [13]. To allow for G4 folding, 40 µM synthetic DNA oligonucleotides (Sigma Aldrich) with 100 µM Tris buffer (pH 7.8) and 100 µM KCl were heated to 90°C for 5 mins, before cooling to room temperature at a rate of 1°C/min. Oligos were mixed with 40 µM ThT and incubated at RT for 5 mins. 25 µL of each oligonucleotide/ThT mixture was then transferred in triplicate to wells of a black walled 96 well plate (Greiner) containing 25 µL of a serial titration of quarfloxin or CX-5461. The plate was incubated for 30 mins at 37°C in the dark and analysed using a FLUO Star Omega Plate reader (BMG Labtech) at Ex 420nm Em 480 nm.

### G4 agarose gel

10 µM oligos were folded as described above and incubated with 100 nM quarfloxin for 30 mins. Samples were mixed with 60% v/v glycerol and loaded into a 1% TAE agarose gel, either mixed with 1X SYBR safe DNA dye (Invitrogen) or without dye and electrophoresed at 100 V for 20 mins. Gels were imaged on a QuantStudio 500 imager using 365 nm for quarfloxin and the epi-blue setting for SYBR-safe.

### Immunofluorescence

Parasites were smeared onto glass microscope slides and fixed for 15 mins in 4% PFA in PBS. Slides were rinsed in PBS, permeabilised for 5 mins in 0.1% Triton-x 100 PBS, and then blocked for 1 h in blocking solution (3% w/v BSA in PBS). Mouse-BG4 antibody (Ab00174-1.1, AbsoluteAntibody) was diluted 1:250 in blocking solution and incubated overnight at 4°C. Slides were rinsed 3 times in PBST and incubated in 1:1000 goat anti mouse Alexa fluor 546 for 1 h, then rinsed again and counterstained with 5 µg/mL DAPI. Slides were rinsed in dH_2_O then mounted with Diamond Prolong antifade (Invitrogen). Samples were imaged on a Zeiss LSM700 confocal, 60X, pinhole – 1 AU, 16 bit at 405 nm for DAPI and 532 nm for BG4 to generate representative images, and by Nikon Microphot SA microscopy (as above for TUNEL assay) using a 100X objective to generate signal intensity measurements from large numbers of cells. Signal intensity was calculated using ImageJ software.

### Flow Cytometry

The published BG-Flow protocol for mammalian cells [49] was adapted for *Plasmodium* parasites. All centrifugation steps were performed at 500 x g for 3 mins using low brake. Synchronous parasites were fixed in freshly made 4% PFA (w/v PBS) containing 0.0025% (v/v) glutaraldehyde in PBS for 30 mins, rinsed for 5 mins in PBS, then permeabilised in 0.05% PBS Tritonx100 (v/v) for 15 mins. Samples were rinsed for 5 mins in PBS then blocked in 3% BSA for 1 h at RT with rotation. Samples were incubated overnight at 4°C with rotation in 1:250 BG4, followed by 1 h 1:1000 donkey anti-mouse IRDye 680RD (LI-COR). Samples were rinsed 3 x 5 mins in PBS following each antibody change and costained in 5 µg/mL DAPI. Parasites were read on an Attune NxT flow cytometer. DAPI signal was detected by 405 nm Violet laser (bandpass 440/50 nm) and BG4 by 637 nm Red (bandpass 720/30 nm) optical filters. For each sample 20,000 events were recorded, doublet discrimination performed by plotting forward scatter height versus area, and singlet population gated for +/+ DAPI/BG4 signal.

### RNA-Seq

Amplification free RNA-Seq was performed as a modified DAFTseq protocol [51]. Briefly, tightly synchronous trophozoites at 30-32 h.p.i were cultured with 2X EC_50_ of either quarfloxin, CX-5461, or no drug for 4 h. Parasites were saponin released and stored in Trizol reagent (Invitrogen). Total RNA was extracted as described previously [67]. Genomic DNA was removed using Turbo DNase clean-up kit (ThermoFisher) according to manufacturer’s instructions and contamination with residual gDNA was checked by PCR amplification using primers designed across an intron of gene PF3D7_0708800 (Supplementary table 4). polyA mRNA was enriched by using NEB polyA module according to kit instructions. First strand cDNA was synthesised using Superscript IV kit (ThermoFisher). Each 20 µL reaction contained 5X FSS buffer, 0.4 µg/µl Oligo d(T) primers, 0.4 µg/µl random primers, 10 mM dNTPs, 0.5 µg/µl ActinomycinD, 0.1 M DTT, 1 µL Superscript IV reverse transcriptase, 20 U RNaseOUT, 9.5 µL mRNA. Samples were subjected to 25*°*C for 10 min, 42*°*C for 60 min, 4*°*C hold. cDNA was cleaned using Ampure XP beads. Second strand cDNA synthesis was performed using NEB Next Ultra II Second Strand Synthesis Module. Each reaction was performed in 50 µL containing 5 µL 10X second strand buffer and 4 µL enzyme mix. Synthesis was performed in a thermocycler at 16*°*C for 2.5 h and a second Ampure bead clean-up was performed. Following quality control check of cDNA, samples were then processed and sequenced at the Cambridge Genomic Services sequencing facility. Briefly, samples were sheared using a Covaris M220, libraries were prepared using KAPA HyperPrep kit (Roche) and samples were sequenced using Illumina NextSeq 550 MO. FASTQ files were accessed from Ilumina BaseSpace and processed for alignment.

### RNA-Seq analysis

Paired end reads from FASTQ files were trimmed using Trimmomatic (v0.39) [68,69] and aligned to *P. falciparum* 3D7 genome v58 [70], using HISAT2 (v2.2.0) [71] and were indexed using BAMtools [72]. Reads were counted using featureCounts from the SubRead (v2.0.2) package [73]. Differential expression was performed in RStudio with R (3.6.3) [74,75] using DESeq2 (v1.36) with adjusted p value <0.05 and log2FC >1 [76]. Heatmaps and volcano plots were used to visualise data spread, and assess for noise across samples, following which outlier data was removed (Supplementary Figure 7). Multiple R packages were used to manage and visualise data [77–79]. Gene ontology enrichment was performed using ReviGO [52].

### Validation of RNA-Seq by qRT-PCR

Follow-up validation was performed using primers designed to 6 differentially expressed genes as determined by RNA-Seq. Parasites were drug treated as above, extracted by phenol-chloroform, and cDNA synthesised using SensiFAST cDNA synthesis kit (Bioline). Reactions were performed in a QuantStudio6 pro real-time PCR system (Applied Biosystems) using SensiFAST SYBR Lo-ROX kit (Bioline). Melt curve analysis was performed. Mean Ct values were calculated from 3 biological repeats and normalised against housekeeping genes PF3D7_0717700 (serine-tRNA ligase) and PF3D7_1444800 (aldolase) to find ΔΔCt. Fold change against the untreated control was calculated and plotted.

### Analysis and statistical tests

Analysis of microscopy images and densitometry were performed in ImageJ unless otherwise stated [80]. Statistical analysis was performed in GraphPad Prism v8.

## Acknowledgements

We thank Prof. Karl Hoffmann (Aberystwyth University) for a generous gift of Quarfloxin. We are especially grateful to Mr Anders Jenson for preliminary experimental work, Dr Francis Totanes for assisting with RNA-Seq experiments and data handling, Dr Anna Protasio and Dr Lia Chappell (Sanger Institute) for advice on RNA-Seq experiments, and Dr Katrin Paeshke for sharing the BG4 flow cytometry protocol ahead of publication. We recognise the knowledge and expertise of Dr Joana Cerveira, Dr David Carpentier and Dr Julien Bauer of the Pathology Department’s flow cytometry, microscopy and sequencing facilities respectively.

## Supplementary materials

**Supplementary Figure 1.**
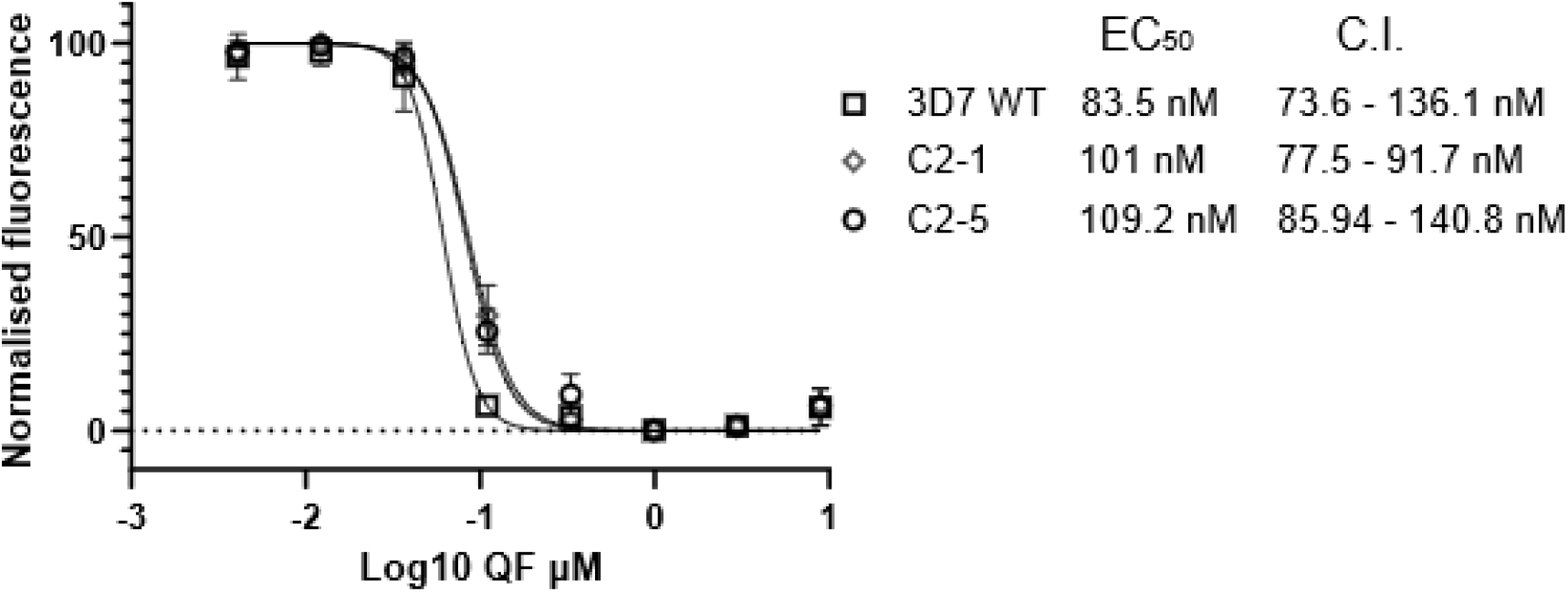
Quarfloxin is ‘irresistible’ in culture. P. falciparum 3D7 parasites were cultured with increasing concentrations of quarfloxin for several months, until stable growth was achieved. This was possible only up to ∼2X EC_50_, with further increases causing complete death of the culture. Cultures were then clonally diluted (still in quarfloxin) prior to MSF analysis. Data from two clones, C2-1 and C2-5, are shown here, alongside the untreated 3D7 strain.

**Supplementary Figure 2.**
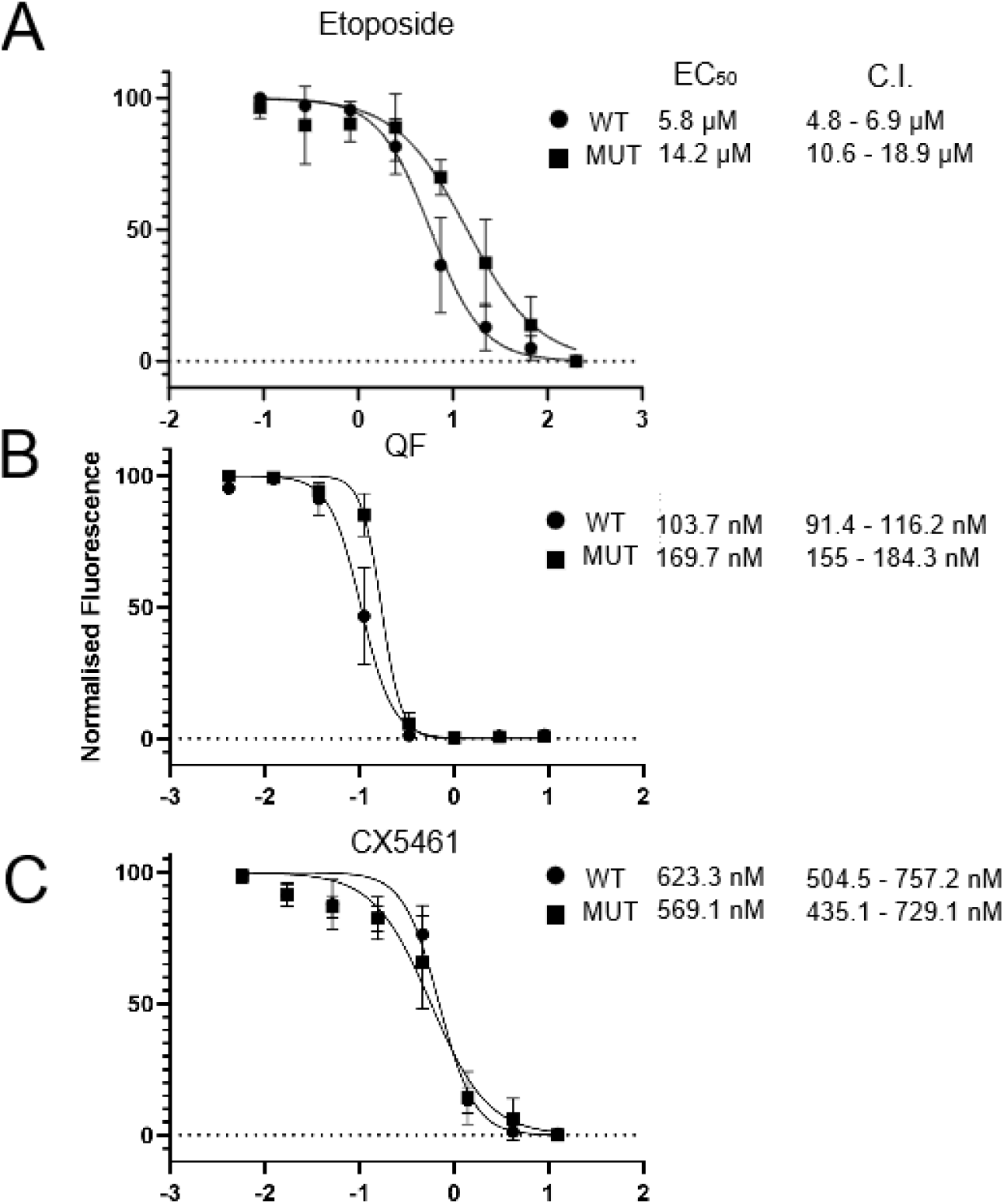
Quarfloxin and CX5461 do not primarily act through topoisomerase interaction. Dose response curve for three drugs on matched parasite lines carrying a Pf RAD54 gene that is either wildtype (WT) or mutated (MUT) at the rG4 locus. The G4MUT line contains increased levels of RAD54 protein, increasing parasite tolerance to A) etoposide, a topoisomerase targeting compound, by ∼2.5-fold. By contrast, the EC_50_ values on these lines for B) quarfloxin differed by ∼1.5-fold and C) CX-5461 did not differ.

**Supplementary Figure 3.**
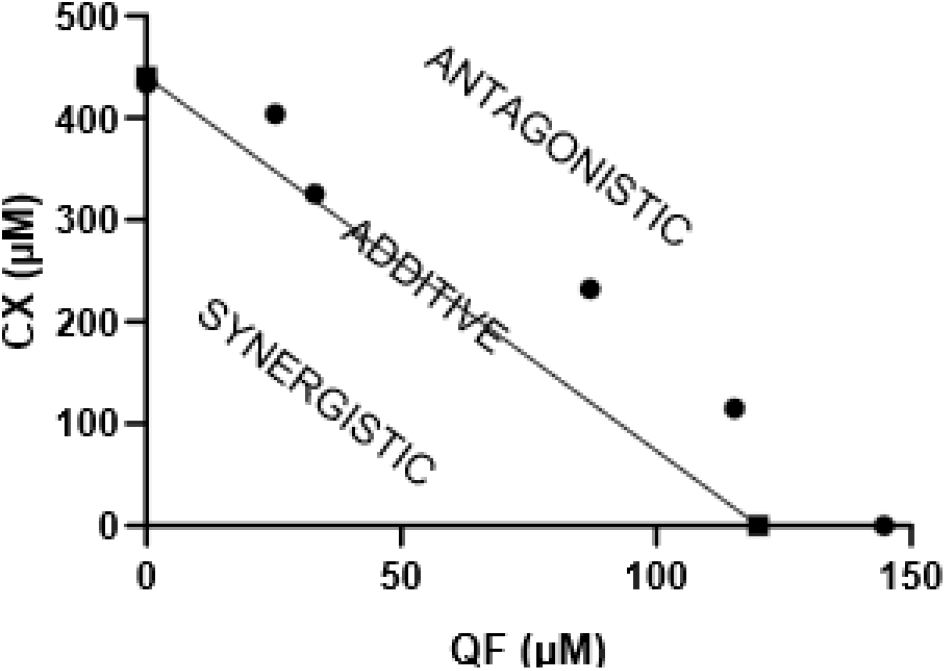
Quarfloxin and CX-5461 do not synergise as co-treatments. Isobologram analysis of the interaction between quarfloxin and CX-5461. EC_50_ values of quarfloxin and CX-5461 alone were plotted to find the additive line. Mixed ratios of quarfloxin and CX-5461 were analysed by MSF and EC_50_ values plotted. Values fall generally above the additive line, and therefore the compounds are not considered synergistic but may be weakly antagonistic.

**Supplementary Figure 4.**
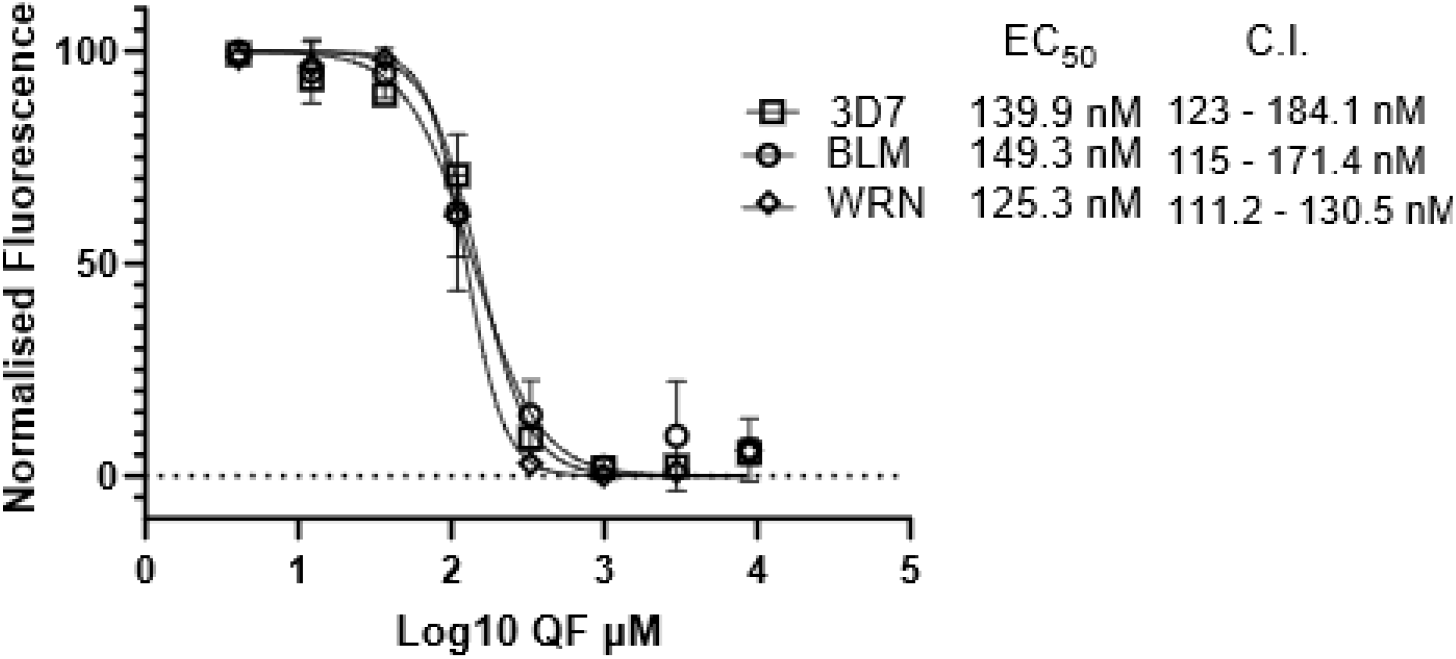
Quarfloxin is not significantly more toxic to parasites mutated in the RecQ helicases PfBLM and PfWRN. Dose responses to quarfloxin were measured in parasite lines mutated in the G4-resolving helicases BLM and WRN. The sensitivities of these lines did not differ significantly from wild-type parasites.

**Supplementary Figure 5.**
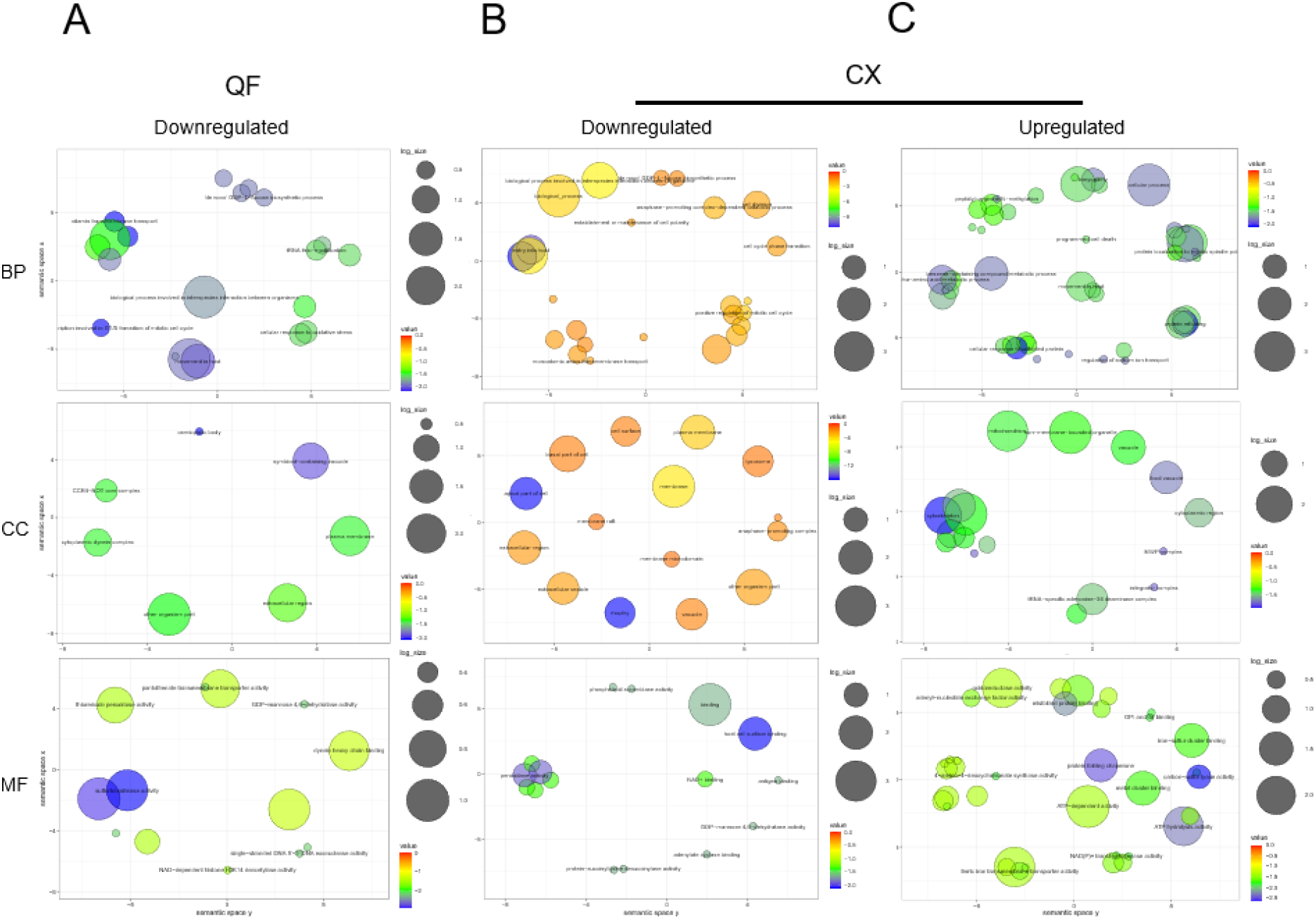
GO enrichment analysis of DE genes with ≥2-fold changes. Biological process (BP), cellular compartment (CC) and molecular function (MF) GO term enrichment was performed for A) quarfloxin downregulated DE genes, B) CX-5461 downregulated and C) CX-5461 upregulated DE genes. Significant terms were plotted in R, P < 0.05.

**Supplementary Figure 6.**
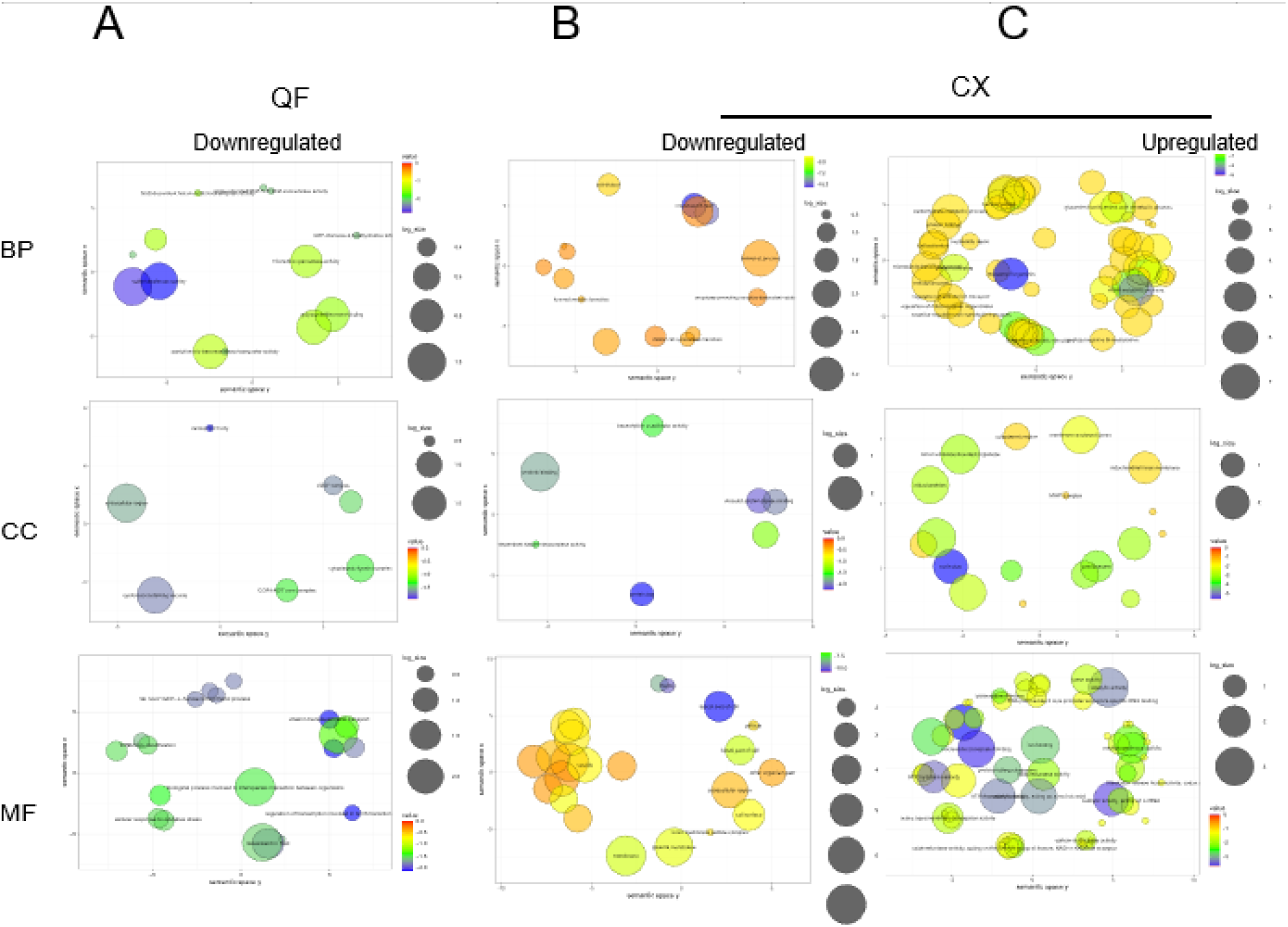
GO enrichment analysis of DE genes with ≥1.5x fold changes. Fold change threshold was reduced to 1.5 and enrichment was repeated as in Supplementary Figure 5. Significant terms were plotted in R, P<0.05.

**Supplementary Figure 7.**
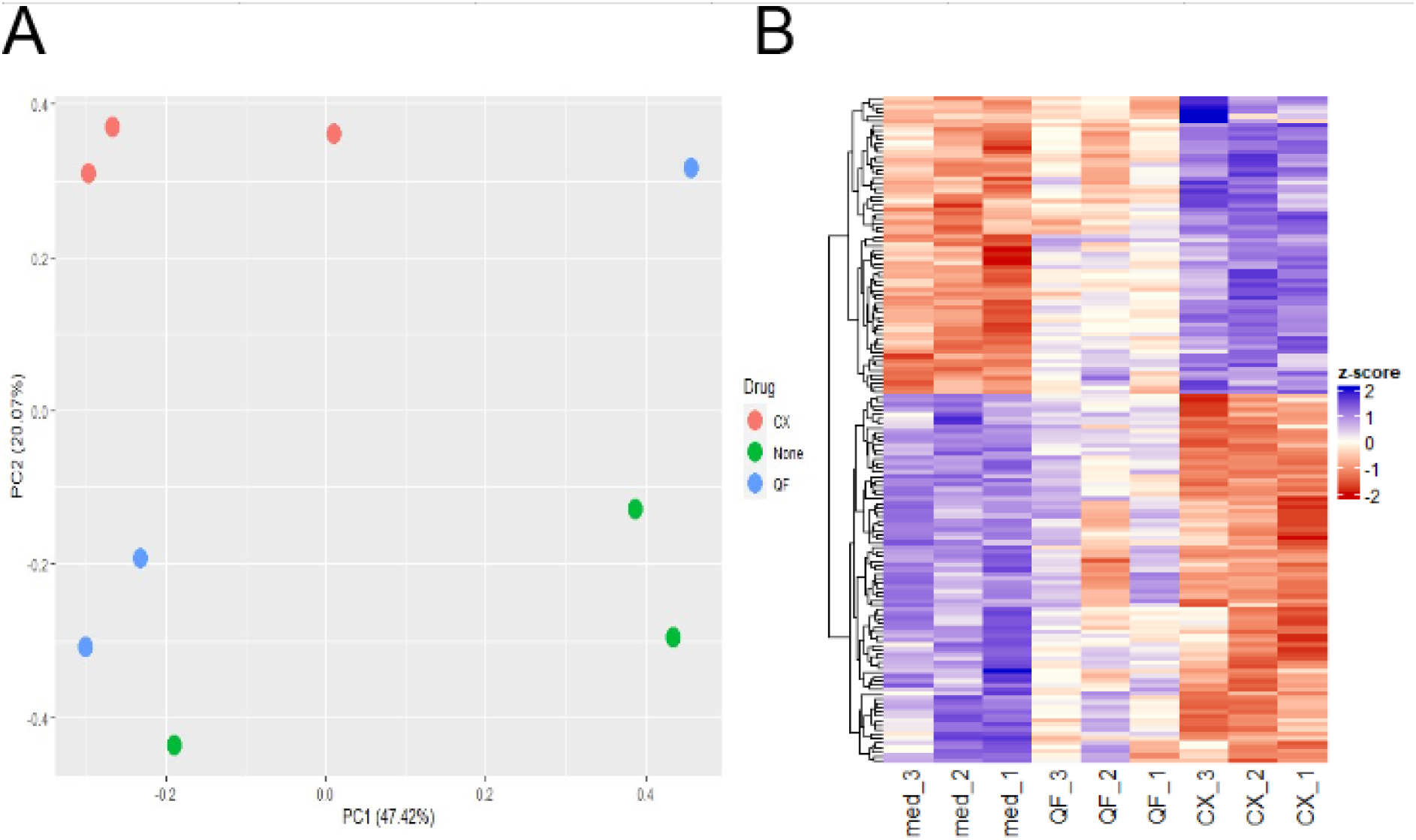
Spread of DE data between and within samples. A) PCA plots show clustering of individual biological repeats (3 repeats per treatment type). B) Heatmap of top 150 DE genes as determined by FDR. Sample QF2 was removed as outlier.

Supplementary Table 1. Differentially expressed genes in quarfloxin-treated parasites vs no drug control, adj. P<0.05, log2FC >1.

Supplementary Table 2. Differentially expressed genes in CX-5461-treated parasites vs no drug control, adj P<0.05, log2FC>1.

Supplementary Table 3. Differentially expressed genes that were found to contain G4s in previously published datasets. Datasets used include G4hunter prediction (scoring threshold >1.2) and rG4seq with and without PDS [9, 12].

Supplementary Table 4. Oligos used in all experiments and their sequences.

